# Elevated *de novo* fatty acid biosynthesis gene expression promotes melanoma cell survival and drug resistance

**DOI:** 10.1101/441303

**Authors:** Su Wu, Anders M. Näär

## Abstract

Elevated *de novo* fatty acid biosynthesis (DNFA) is a hallmark adaptation in many cancers that supports survival, proliferation, and metastasis. Here we elucidate previously unexplored aspects of transcription regulation and clinical relevance of DNFA in melanomas. We show that elevated expression of DNFA genes is characteristic of many tumor types and correlates with poor prognosis. Elevated DNFA gene expression depends on transcription factor SREBP1 in multiple melanoma cell lines. SREBP1 predominantly binds to the transcription start sites of DNFA genes, directly regulating transcription via RNA polymerase II recruitment and productive elongation. We find that SREBP1-regulated DNFA represents an intrinsic survival mechanism in melanoma cells, regardless of proliferative state and oncogenic mutation status. Indeed, malignant melanoma cells exhibit elevated DNFA gene expression after pro-survival signaling pathways are blocked (e.g. by the BRAF inhibitor vemurafenib). Altogether, these results implicate SREBP1 and DNFA enzymes as enticing therapeutic targets in melanomas.

## Introduction

Cancer cells characteristically achieve hallmark traits that facilitate proliferation, survival, and metastasis^1–3^. One hallmark adaptation is *de novo* fatty acid synthesis (DNFA), metabolic conversion of carbohydrates into lipids *via* acetyl-CoA and NADPH with the aid of multiple lipogenic enzymes, including ATP citrate lyase (ACLY), acyl-coenzyme A synthetase 2 (ACSS2), acetyl-CoA carboxylase (ACACA), fatty acid synthase (FASN), and stearoyl-CoA desaturase (SCD)^4^. DNFA occurs in cancer cells and certain types of healthy cells^5^. In liver cells, DNFA pathway activity is regulated at the level of DNFA enzyme gene expression in response to dietary lipids (e.g. polyunsaturated fatty acids^6–8^), and hormonal cues such as insulin^9^. DNFA is also increased in normal cells and tissues during embryonic development and adipogenesis to satisfy elevated lipid demands during cell proliferation and fat storage processes, respectively^10,11^.

The transcription factor Sterol Regulator Element-Binding Protein 1 (SREBP1) plays a central role in controlling DNFA gene expression, and thus serves as a master regulator of cellular FA/lipid production^12,13^. There are two major mechanisms involved in SREBP1 regulation: mRNA expression and proteolytic processing^14^. The *SREBF1* gene, whose transcription is regulated by insulin and cellular/membrane lipids, encodes a SREBP1 precursor protein embedded in the endoplasmic reticulum membrane through two transmembrane domains^15–17^. In response to depletion of cellular and membrane lipids, SREBP cleavage-activating protein (SCAP) escorts SREBP1 to the Golgi apparatus, where its active, nuclear form (nSREBP1) is released by Site 1 and Site 2 proteases^18–20^, allowing nuclear translocation and binding to the promoters of target genes. nSREBP1 activates the transcription of DNFA genes, in concert with other transcription factors LXR^21^, USF1^22^, NFY1^23^ and SP1^24^, and co-activators MED15^24^ and CREBBP^25^. nSREBP1 also participates in activation of *SREBF1* mRNA expression by binding to its own promoter^25^, thus the levels of DNFA mRNAs parallel the changes in *SREBF1* expression^12^.

Elevated DNFA has been demonstrated in many tumor types^26^, including breast cancer^27^, prostate cancer^28^, and glioblastoma multiforme^29^. The prevailing thought is that hallmark traits, such as DNFA, arise chiefly as a consequence of pro-survival signaling pathways driven by genetic alterations of oncogenes and tumor suppressors^30–33^. Supposed dependence of tumor cells on a single oncogenic driver or pathway to sustain proliferation and/or survival has guided the development of targeted cancer therapies^34,35^. However, in clinical settings, tumors harbor highly diverse genetic alterations, and exhibit stochastic evolution^36^. The prognostic and therapeutic value of driver alterations is therefore frequently limited^37–40^. Resistance to targeted therapies is common in tumors due to mechanisms related to reactivation or bypass of downstream signaling pathways^41^. It is unclear whether oncogene alterations maintain hallmark traits such as DNFA in malignant tumors. Furthermore, potential interaction between oncogenic drivers and DNFA has not been fully investigated, especially under the selection pressure of targeted therapies.

We show here that elevated expression of SREBP1 target genes (e.g. key DNFA enzymes such as SCD) is significantly associated with poor prognosis in cancers, including melanomas. Our detailed mechanistic analyses reveal that SREBP1 and DNFA play crucial roles in melanoma cell proliferation and survival, as well as resistance to targeted therapies (e.g., the BRAF inhibitor vemurafenib) in melanoma cancer cells.

## Results

### Elevated mRNA expression of DNFA enzymes is prevalent in many cancers, including malignant melanomas, and has prognostic value

Elevated lipogenic enzyme activities have been reported in colon, breast and prostate cancers^42–44^. We analyzed the expression of mRNAs encoding the DNFA enzymes SCD, FASN, ACLY and ACSS2 using RNA-Seq data from 30 diverse cancer types in The Cancer Genome Atlas (TCGA) (Fig. 1a, b and Supplementary Fig. 1a, b). We found that DNFA enzyme expression varies widely among different types of cancers, with melanoma exhibiting among the highest levels of expression. Kaplan-Meier analyses indicate that elevated *SCD*, *FASN* and *ACLY* mRNA expression each correlates with poor prognosis in the skin cancer study group (SKCM) (Fig. 1d, and Supplementary Fig. 1d, f). Expression of *SCD* and *FASN* likewise was significantly elevated in all cancers considered collectively (Fig. 1c and Supplementary Fig. 1c), whereas for consideration of SKCM as distinct among cancer types — *ACLY* was not (Supplementary Fig. 1e). Prior reports have described activation of enzymes for both DNFA and cholesterol biosynthesis triggered by oncogenic pathways^45,46^. However, we observed low expression of *HMGCS1* and *HMGCR*, rate-limiting enzymes in the cholesterol biosynthesis pathway^47^, in melanomas compared to other cancer groups and no prognostic value for their expression (data not shown). Among healthy tissues, skin *SCD* expression is low (Supplementary Fig. 2a); yet among tumor tissues, skin *SCD* expression is elevated (Fig. 1a). This contrasts with other cancers such as those derived from the liver, which exhibits relatively high *SCD* expression among healthy (behind only brain and adipose, Supplementary Fig. 2a) and tumor tissues (Fig. 1a) alike.

**Figure 1.**
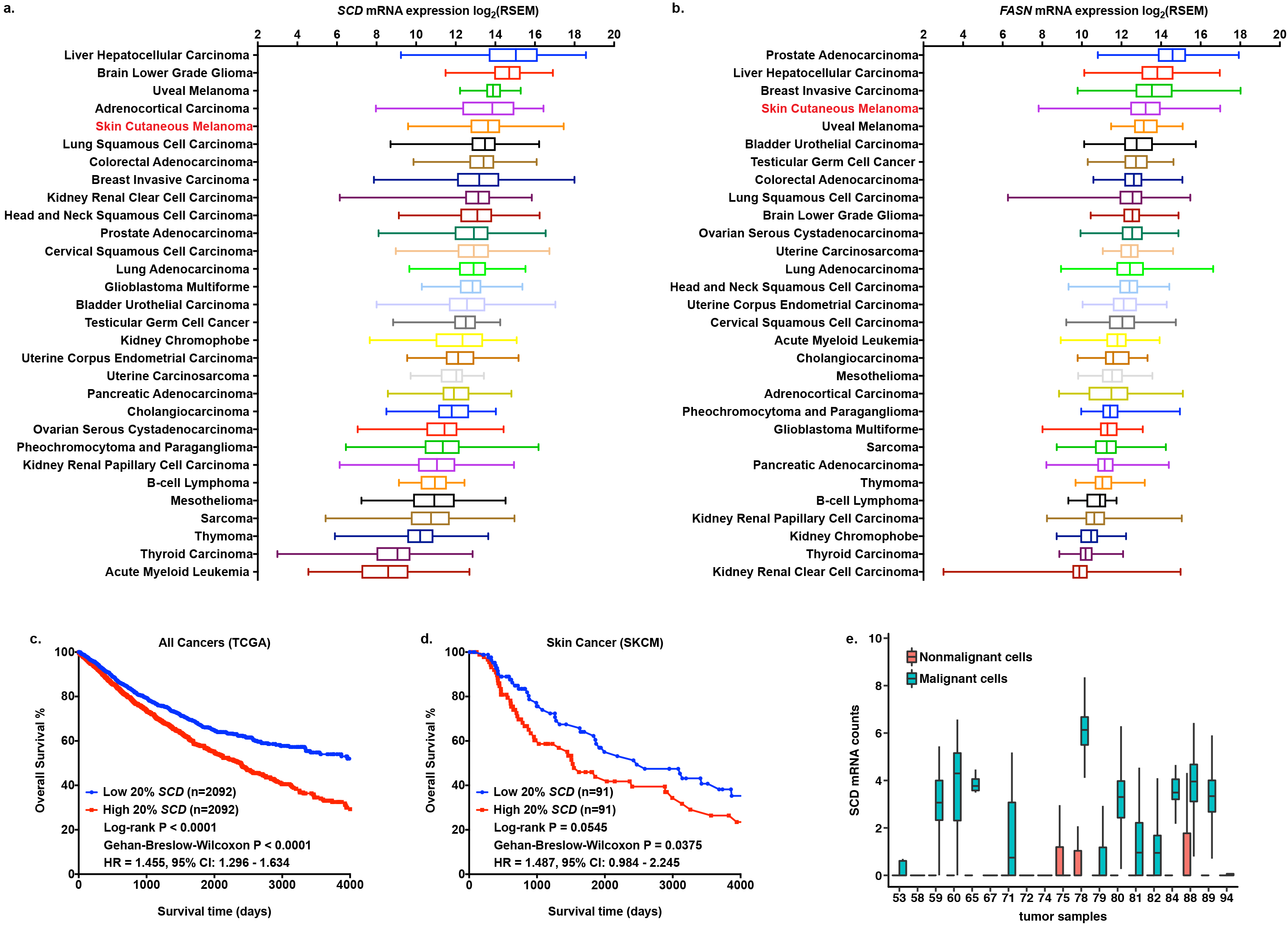
Elevated expression of DNFA genes is prevalent in many cancers, including melanomas, and has prognostic value. **a**, **b**, Expression of *SCD* and *FASN* genes was compared using RSEM normalized RNA-Seq data from 10,210 tumor samples downloaded from The Cancer Genome Atlas (TCGA). The box and whisker plots represent gene expression in 30 TCGA cancer types. **c**, **d**, Differences in overall survival rates are computed between patients with top 20% *SCD* RNA-Seq counts in their tumor samples and those with bottom 20% *SCD* expression, as the Kaplan-Meier plot in all cancer patients and skin cancer patients (SKCM) from TCGA dataset. **e**, Boxplot shows average mRNA reads of *SCD* in 4,645 single cells from tumor samples of 19 melanoma patients (GSE72056). *SCD* expression was compared between malignant and nonmalignant cells.

To exclude the possibility that *SCD* overexpression in melanomas is due to tissue-specific upregulation, we compared *SCD* expression using normalized RNA-Seq data from the skin cancer group in TCGA and normal skin tissue group in the Genotype-Tissue Expression (GTEx) database. *SCD* expression is much higher in the cancer group versus normal skin group (Supplementary Fig. 2b). Moreover, principal component analysis (PCA) shows that skin tumors are distinct from samples of normal skin, based on comparison of expression levels of five DNFA genes (Supplementary Fig. 2c). To confirm that DNFA enzyme expression was specific to malignant cells within bulk tumors, we further analyzed single cell RNA-Seq data from melanoma patient samples^48^. High expression of DNFA enzyme genes such as *SCD*, *FASN*, and *ACACA* was confined to malignant cells, with very low expression in healthy adjacent tissue (Fig. 1e and Supplementary Fig. 2d-g). *BRAF* and *NRAS* mutations, while being well-known as risk factors and drivers of cancer onset, have limited prognostic significance for overall survival of melanoma patients^49^. Consistently, we observed no significant correlation between DNFA enzyme expression and common oncogenic driver mutations in SKCM from TCGA (Supplementary Fig. 3a-l).

### SREBP1 controls highly active DNFA gene expression in melanoma cells

To test whether SREBP1 drives elevated DNFA enzyme expression in melanoma cells, we depleted the mRNA encoding SREBP1 (*SREBF1*) with antisense oligonucleotides (ASOs) and siRNAs. We found that depletion of *SREBF1* with both siRNA and ASO agents was accompanied by decreasing protein levels of SREBP1 and DNFA enzymes (Fig. 2a). We confirmed that pooled siRNA and ASO agents effectively depleted SREBP1 protein, both the cytoplasmic precursor form and the mature form in the nucleus (Fig. 2b, c). Among six tested ASOs targeting *SREBF1*, we found that ASO-1 and ASO-4 yielded strong inhibition of SREBP1 protein and DNFA enzyme production in melanoma cells (Fig. 2a, b). ASO-1 and ASO-4 are more potent than single or pooled siRNAs for *SREBF1*, as 5 nM of ASOs achieved a similar degree of inhibition on DNFA enzyme production as 50 nM of siRNAs (Fig. 2a and Supplementary Fig. 4b-f), even when siRNAs decreased the level of *SREBF1* mRNA more than ASOs (Supplementary Fig. 4a). Our interpretation is that, other than mRNA degradation mechanism similar to siRNAs, ASOs may engage in steric translation inhibition of the *SREBF1* mRNA^50^. We found that ASO-4 inhibits DNFA gene expression commensurately with dosage (Supplementary Fig. 4g). We also observed a dose-response relationship of diminished cell viability with increased ASO concentration that was not evident with siRNAs (data not shown).

**Figure 2.**
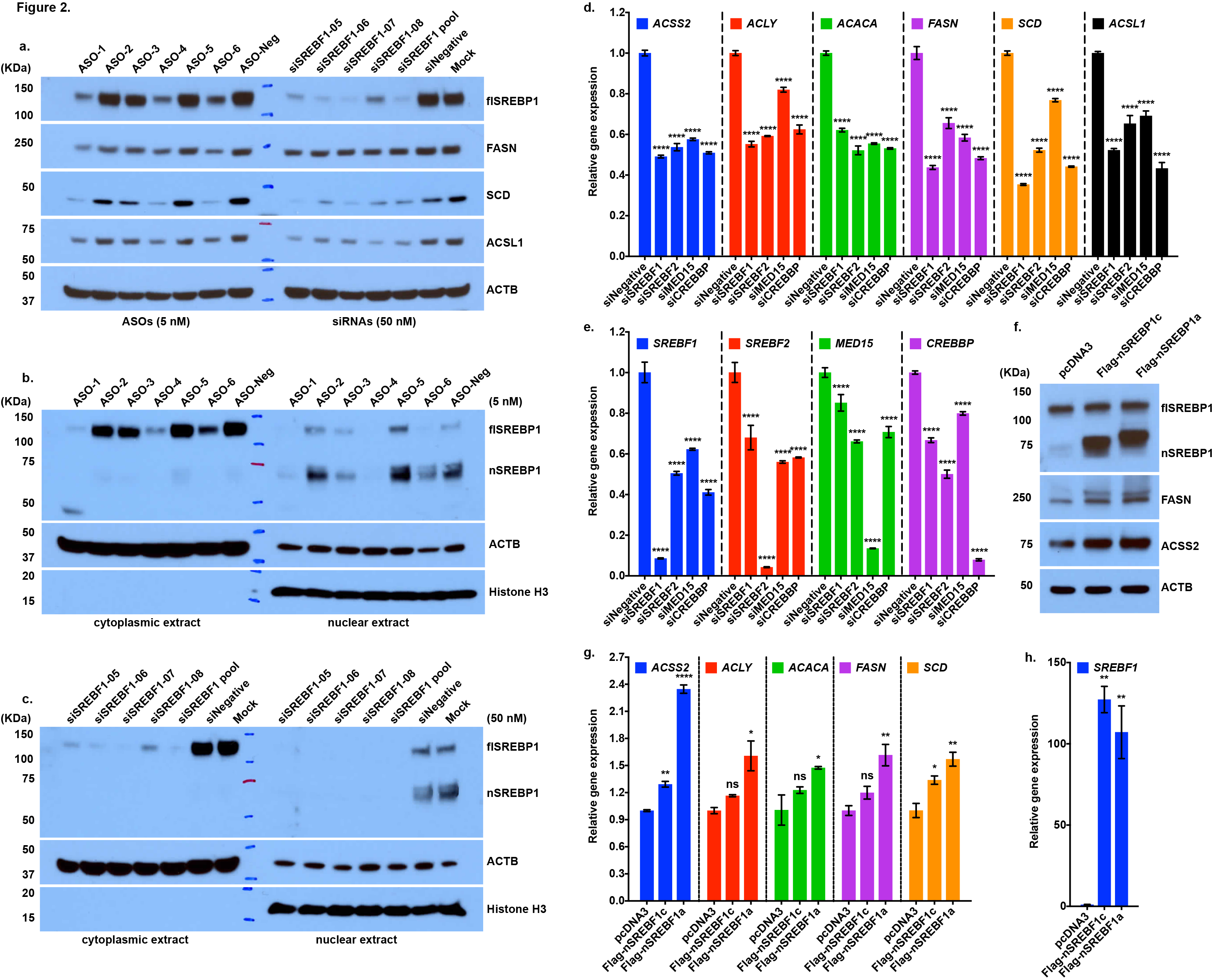
SREBP1 regulates the elevated DNFA gene expression in melanoma cells. **a**, HT-144 cells were treated with ASOs, individual siRNA agents or pooled siRNAs (all individual agents combined) to deplete SREBP1 in 1% ITS medium. Total cell lysates were assayed with immunoblot by the indicated antibodies. HT-144 cells were transfected with (**b**) ASOs, (**c**) individual siRNA agents or pooled siRNAs in 1% ITS medium. Nuclear and cytoplasmic extracts were isolated for Western blot analysis for full length (fl) and nuclear (n) SREBP1 protein after treatment. **d**, **e**, HT-144 cells were transfected with the pooled siRNAs (50 nM) in 1% ITS medium for three days to deplete *SREBF1*, *SREBF2*, *MED15* or *CREBBP.* RT-qPCR assay of mRNA shows relative expression of DNFA enzymes from siRNAs treatment groups to that of negative control siRNA treatment (siNegative). **f-h**, HT-144 cells were transfected with plasmids carrying the transcriptionally active N-terminal portion of SREBP1a (nSREBP1a), N-terminal portion of SREBP1c (nSREBP1c) or empty vector (pcDNA3) for two days. **f**, the total cell lysate was analyzed by Western blot assay using the indicated antibodies. **g**, **h**, the mRNAs were analyzed with RT-qPCR assay. The bar graphs show the relative expression of DNFA enzymes to that of pcDNA3 (control) transfection group. Data are expressed as mean ± SD and quantified from triplicates. One-way ANOVA tests were performed. ns, not significant; *, P < 0.05; **, P < 0.01; ***, P < 0.001; ****, P < 0.0001.

To investigate various potential activators of DNFA enzyme expression, we depleted *SREBF1*, *SREBF2*, and co-activators *MED15* and *CREBBP* with siRNAs, and examined expression of DNFA enzymes across melanoma cell lines HT-144 (Fig. 2d-e), A375 (Supplementary Fig. 4i-j) and MEL-JUSO (Supplementary Fig. 5a-b). We observed a similar range of mRNA reductions (50%–70%) for most DNFA enzymes after *SREBF1* depletion, and lesser reduction after co-activator depletion in the three melanoma cell lines (Fig. 2d, Supplementary Fig. 4i and Supplementary Fig. 5a). Depletion of co-activators *MED15* and *CREBBP* individually or together has some impact on DNFA gene expression, but to a lesser extent than depletion of *SREBF1* (Supplementary Fig. 4i and Supplementary Fig. 5a). *SREBF2* depletion is more specific for genes in cholesterol biosynthesis (Supplementary Fig. 4h and Supplementary Fig. 4k), in line with previous studies in liver^51^. *SREBF2* also affected DNFA enzyme expression, especially in combination with *SREBF1* depletion (Supplementary Fig. 5a and Supplementary Fig. 5c-d. The role of *SREBF2* in regulation of DNFA enzyme expression may be transitive via *SREBF1*^52^, since *SREBF1* expression decreased after *SREBF2* depletion (Supplementary Fig. 5e), or it may be acting in a partially redundant manner^53^. Thus, we confirmed the known transcription regulatory role of SREBP1 in controlling DNFA gene expression in multiple melanoma cell lines.

To understand the dynamics of DNFA gene expression, we performed a time course study of DNFA expression in A375 cells cultured under SREBP1-activating conditions (1% ITS medium); as expected, *FASN* and *SCD* displayed increased expression at consecutive time points under these conditions (Supplementary Fig. 5c-f). *SREBF1* depletion by siRNA decreased DNFA gene activation in A375 cells on days 3 and 5, in agreement with the notion that SREBP1 directly regulates expression of DNFA enzyme genes. Next, we evaluated the response to SREBP1 over-expression using transfected plasmids encoding the constitutively active (nuclear) form of SREBP1a (nSREBF1a), nuclear form of SREBP1c (nSREBF1c) or control vector in HT-144 melanoma cells. We observed elevated expression of DNFA enzyme proteins by Western blotting (Fig. 2f) and DNFA genes by RT-qPCR analysis (Fig. 2g-h) in response to increased expression of both SREBP1a and SREBP1c isoforms. SREBP1c is the predominant isoform in adult organs such as liver and adipose tissues^54^, while SREBP1a is more abundant in proliferating embryonic cells and in cancers. With a longer transactivation domain, SREBP1a interacts more avidly with co-activators and thus exhibits stronger transcription activity than SREBP1c^24,55^. Consistently, we observed higher expression of DNFA genes after overexpressing SREBP1a than SREBP1c. Our overall interpretation of the results is that SREBP1 is sufficient for highly active DNFA gene expression in melanoma cells, which is in accord with previous published studies in other cancer cell types.

### SREBP1 maintains highly active DNFA gene expression through RNAP II recruitment and productive transcription elongation

To further assess the gene regulatory function of SREBP1 in melanoma cells, we carried out RNA-Seq analysis after *SREBF1* depletion with pooled siRNAs and individual ASOs in HT-144 cells, followed by principal component analysis (PCA) on RNA-Seq data to characterize the gene expression patterns after *SREBF1* depletion. The two principal components in the PCA biplot represent over 70% of the overall gene expression changes (Fig. 3a and Supplementary Fig. 6a). *SREBF1* and *SCD* are among the top six contributors to data separation and they align with principal component 2 (PC2) (Fig. 3a). We found that the major contributors to PC2 were *SREBF1* and DNFA genes including *FASN* and *SCD* (Supplementary Fig. 6b). *SREBF1* siRNA, ASO-1 and ASO-4 obtained similar vertical separations to all negative controls in PC2, and they all reduced mRNA reads of *FASN* and *SCD* compared to negative controls (Supplementary Fig. 6e-g). These results indicate that the most profound effects after *SREBF1* depletion by both ASO and siRNA are on DNFA genes. ASO-1 has significant lateral separation from ASO-4 and *SREBF1* siRNA on principal component 1 (PC1) (Fig. 3a). We found that changes of *SPIRE1* and *USP9X* expression are the major contributors to PC1 (Supplementary Fig. 6c) and are only affected by ASO-1 (Supplementary Fig. 6h, i). This result indicates that ASO-1 has specific off-target effects on *SPIRE1* and *USP9X* genes. Multiple negative controls grouped together, and ASO-4 is close to *SREBF1* siRNA on the PCA biplot, consistent with the result of hierarchical clustering analysis on the same RNA-Seq data (Supplementary Fig. 6d). Hence, ASO-4 has similar specificity as pooled *SREBF1* siRNAs for *SREBF1* depletion.

**Figure 3.**
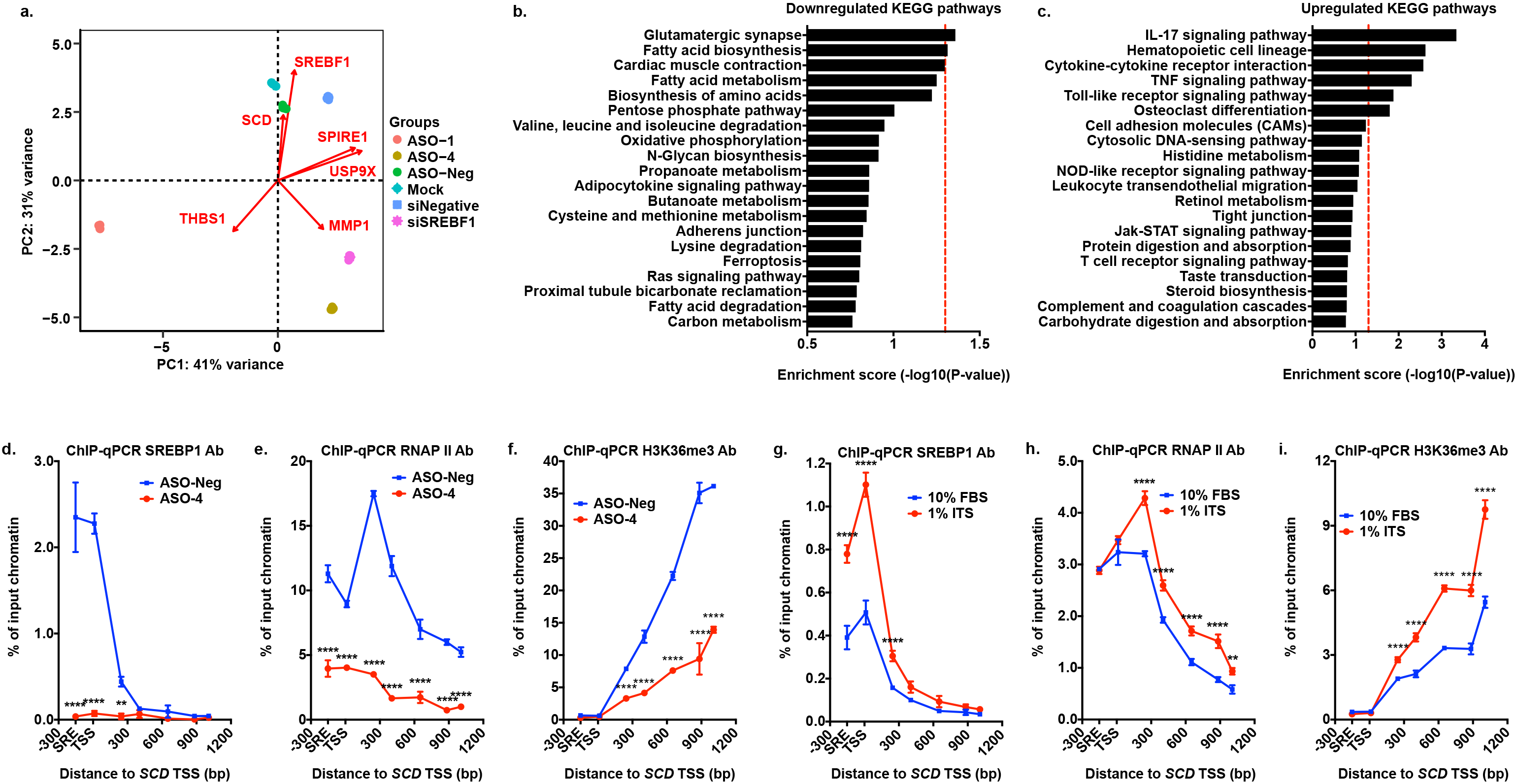
The mechanism by which SREBP1 causes highly active DNFA gene expression in melanoma cells. **a**, mRNAs of HT-144 cells were sequenced after ASOs (5 nM) or pooled siRNAs (50 nM) treatment in 1% ITS medium for three days. RNA-Seq data were analyzed with DESeq2 and principal component analysis (PCA). **b**, **c**, top 20 enriched signaling and metabolic KEGG pathways were discovered from differentially expressed genes (siSREBF1 vs siNegative group) in RNA-Seq analysis using Generally Applicable Gene-set Enrichment (GAGE). Red dash line marks P value = 0.05. **d-f**, HT-144 cells were transfected with ASO-4 (5 nM) or control ASO (5 nM), cultured in 1% ITS medium. Percentage of input DNA was compared between two treatments for the indicated antibodies at the 5’ promoter region of the *SCD* gene. **g-i**, ChIP-qPCR analyses detected DNA pulldown using indicated antibodies at the *SCD* gene in HT-144 cells. ChIP-qPCR signals were compared between cells cultured in 10% FBS and 1% ITS medium condition. Data were presented as mean ± SD and quantified from 3 triplicates. Two-way ANOVA tests were performed. ns, not significant; *, P < 0.05; **, P < 0.01; ***, P < 0.001; ****, P < 0.0001.

To assess the overall transcriptome affected by *SREBF1* depletion, we examined the differentially expressed genes (DEGs) in the RNA-Seq data (siSREBF1 vs siNegative groups) (Supplementary Fig. 6j). Using gene-set enrichment analyses (GSEA) with Kyoto Encyclopedia of Genes and Genomes (KEGG) pathways and Gene ontology (GO) terms on DEGs, we determined that fatty acid biosynthesis pathway and lipid metabolism pathways are enriched among downregulated genes after *SREBF1* depletion (Fig. 3b, and Supplementary Fig. 6k). Cellular inflammatory response pathways were significantly enriched in upregulated genes after *SREBF1* depletion (Fig. 3c, and Supplementary Fig. 6l), including Toll like receptor and tumor necrosis factor (TNF) signaling pathways that mediate tumor cytotoxicity^56^. We suspect that two major altered programs after *SREBF1* depletion — decreased fatty acid biosynthesis activities and enhanced inflammatory response activities — may facilitate cell death in melanomas.

The time course RT-qPCR (Supplementary Fig. 5c, d) and RNA-Seq (Fig. 3a) analyses suggest direct regulation by SREBP1 of *FASN* and *SCD* expression in melanoma cells. To further elucidate the molecular mechanism of DNFA gene activation in melanoma cells, we used chromatin immunoprecipitation (ChIP)-qPCR analysis to detect occupancy of SREBP1 and RNA polymerase II (RNAP II), and a histone mark of transcription elongation (H3K36me3)^57^ on DNFA genes. SREBP1 depletion by ASO-4 treatment diminished SREBP1, RNAP II and H3K36me3 signals at the *SCD* promoter (Fig. 3d-f). We observed similar results (albeit with smaller magnitude) for *FASN* (Supplementary Fig. 7a-c). These ChIP-qPCR results together with the RNA-Seq data in Supplementary Fig. 6e-g suggest that removal of SREBP1 at DNFA promoters inhibits transcription activity and mRNA production.

To define the molecular action of SREBP1 at DNFA gene promoters in melanoma cells, we performed ChIP-qPCR analyses in SREBP1-activating (1% ITS medium, no lipids) and SREBP1-repressing (10% FBS medium, with lipids) conditions. We found that 1% ITS medium dramatically increased SREBP1 occupancy at the transcription start sites (TSS) of *SCD* (Fig. 3g) and *FASN* (Supplementary Fig. 7f) in HT-144 cells. The strong RNAP II binding peaks at TSS of *SCD* and *FASN* in both 10% FBS and 1% ITS culture conditions (Fig. 3h and Supplementary Fig. 7g) indicate promoter-proximal pausing of RNAP II^58^. Furthermore, culturing cells in 1% ITS medium increased the occupancy of actively elongating RNAP II (RNAP II S2P) (Supplementary Fig. 7d, i), but not poised RNAP II (RNAP II S5P)^59^ (Supplementary Fig. 7e, j), at the TSS as well as the gene body of both *SCD* and *FASN* genes. Accordingly, levels of the H3K36me3 histone mark, which is associated with transcription elongation, were increased across both genes in cells cultured in SREBP1-activating medium (Fig. 3i and Supplementary Fig. 7h). These results suggest that SREBP1 binding at promoters near the TSS associates with RNAP II recruitment and productive transcription elongation on DNFA genes.

To further explore the occupancy of SREBP1 on DNFA genes in cancers, we analyzed public ChIP-Seq data for SREBP1 from lung cancer, breast cancer and chronic myeloid leukemia (CML) cell lines. ChIP-Seq peaks primarily localize at the proximal promoter regions around transcription start sites (Supplementary Fig. 8a-b). *De novo* motif sequences identified from SREBP1 ChIP-Seq peaks match the known SREBP1 binding motif (Supplementary Fig. 8c). We determined the overlapping genes among DEGs in our RNA-Seq data and public SREBP1 ChIP-Seq data from A549 and MCF7 cell lines (Supplementary Fig. 8d), because of high DNFA gene expression in lung and breast cancers (Fig. 1a). We reasoned that the overlapping genes are likely regulated directly by SREBP1, and performed functional network analysis on this subset. This analysis revealed genes in the PI3K/AKT pathway and the RNAP II elongation complex, in addition to the expected DNFA pathways (Supplementary Fig. 8e). We then performed GSEA on the overlapping genes using expression changes from the RNA-Seq data. Our results show that lipid metabolism pathways are significantly enriched in the overlapping gene set that is downregulated after SREBP1 depletion (Supplementary Fig. 8f), confirming SREBP1 as a direct activator of DNFA genes. Although inflammatory response pathways were significantly upregulated in DEGs from RNA-Seq analysis, they seem not to be direct targets of SREBP1 (Supplementary Fig. 8g). We suspect downregulation of DNFA pathways may change the homeostasis of cellular fatty acids and exert further impact on inflammatory response pathways as well as cell death^60^.

### Melanoma cells exhibit continued and elevated DNFA gene expression after oncogenic signaling blockade with the BRAF inhibitor vemurafenib

To evaluate whether melanoma cells rely on DNFA for cell proliferation and survival, we cultured melanoma cells in the 1% ITS medium, using insulin as a growth factor to stimulate proliferation while constraining cellular lipid availability to DNFA output. The metastatic melanoma tumor-derived cell lines HT-144 and A375 proliferate in both 10% FBS and 1% ITS media, but remain quiescent in 0% FBS medium (Supplementary Fig. 9a, b). By contrast, the primary melanoma tumor-derived cell line, MEL-JUSO, remains quiescent in both lipid-depleted media (1% ITS and 0% FBS medium) (Supplementary Fig. 9c). We depleted *SREBF1* with ASO-4 in several melanoma cell lines cultured under the three media conditions. We found that ASO-4 decreased viability of proliferative and quiescent cells in all conditions (Supplementary Fig. 9d-i). Comparing the conditions with growth factors (10% FBS and 1% ITS), we find that lipid availability in the medium somewhat decreases cellular sensitivity to ASO-4 inhibition. We reason that, although the cancer cells are able to utilize ambient lipids^61^, DNFA is required for cell survival regardless of external lipid availability.

We next investigated whether activated DNFA contributes to the mechanism of resistance to BRAF inhibitors (BRAFi, e.g. vemurafenib) observed in melanoma cells harboring the oncogenic BRAF mutation *BRAF*^*V600E* 62^. We derived two BRAFi-resistant cell lines: HT-144BR from a vemurafenib-sensitive cell line HT-144, and LOXIMVIBR from a vemurafenib-insensitive cell line LOXIMVI, with prolonged vemurafenib treatment (Fig. 4a, Supplementary Fig. 9j). By testing with ASO-4, we determined that vemurafenib-sensitive, vemurafenib-insensitive and vemurafenib-resistant cell lines all depend on SREBP1 for survival (Fig. 4b and Supplementary Fig. 9k). We found that DNFA gene expression was higher in vemurafenib-resistant cell lines than in vemurafenib-untreated cell lines (Fig. 4c and Supplementary Fig. 9l). To confirm that inhibition of DNFA diminishes cell viability, we then used small molecule inhibitors of DNFA enzymes FASN and SCD, which have been reported to decrease growth of a number of tumor types in preclinical studies^63–65^, in 10% FBS (lipid-containing) medium. We observed decreased cell viability in both HT-144 and HT-144BR cells (Fig. 4d, f). The effect was less potent than *SREBF1* depletion by ASO-4; however, we observed much stronger effects on cell survival when combining two inhibitors together. Bliss independence analysis^66^ confirmed positive synergy between FASN and SCD inhibitors (Fig. 4e, g). DNFA thus appears vital to the vemurafenib resistance mechanism.

**Figure 4.**
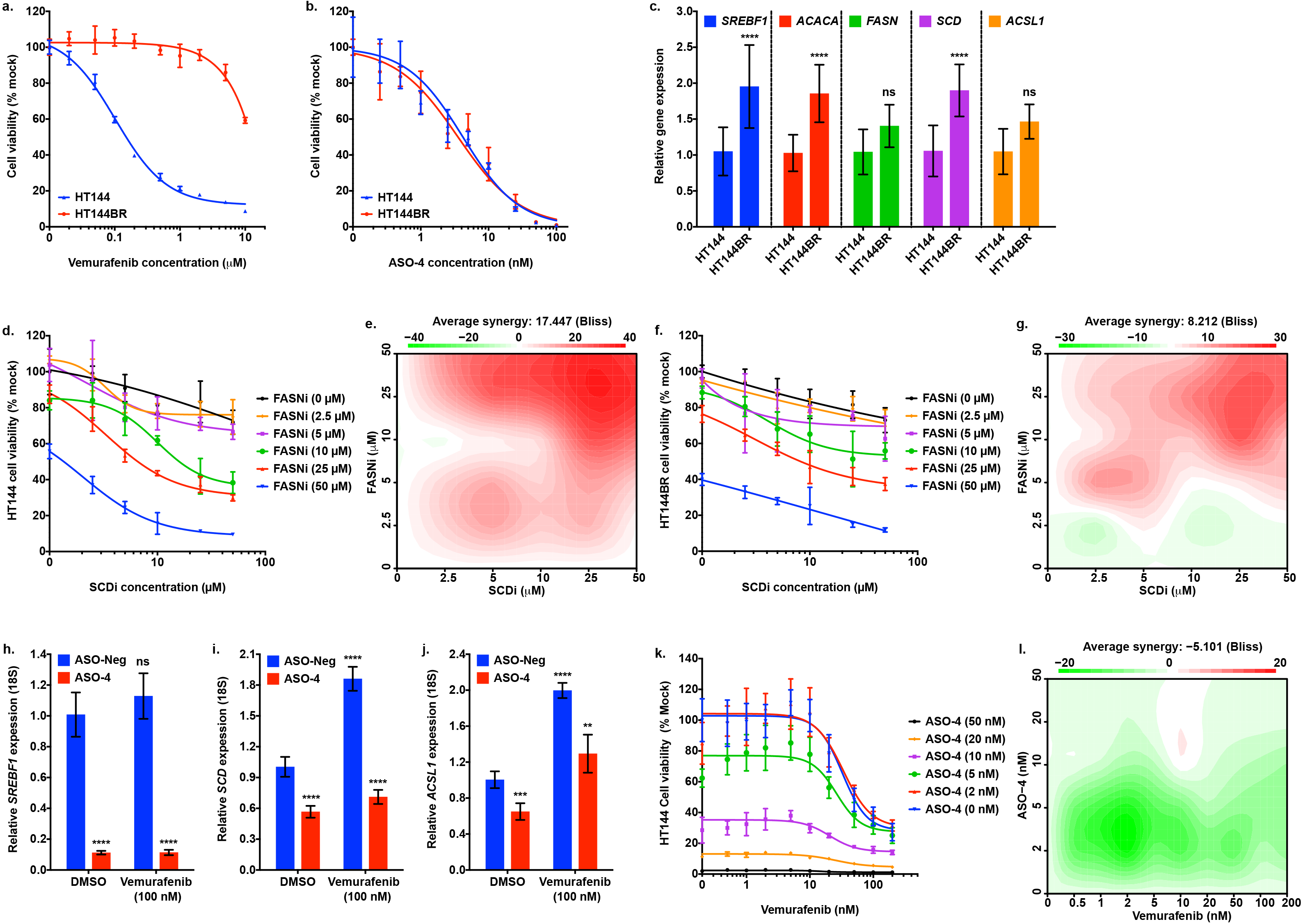
DNFA is necessary for survival and supports drug resistance in melanoma cells. **a-b**, HT-144BR is a BRAF inhibitor resistant HT-144 cell line derived after prolonged vemurafenib treatment (2 μM) in 10% FBS medium for three months. Viability of HT-144 and HT-144BR cells responding to vemurafenib or ASO4 treatment in 10% FBS medium was assessed by cell titer-glo assay. **c**, RT-qPCR assay compared the DNFA gene expression between HT-144 and HT-144BR cells. **d-g**, HT-144 and HT-144BR cells were co-treated with FASN inhibitor (FASNi) GSK 2194069 and SCD inhibitor (SCDi) MF-438 in 10% FBS medium. Cell viability was measured by cell titer-glo assay six days after treatment. The combination responses for two DNFA enzyme inhibitors were evaluated by Bliss independence model. Positive bliss scores indicate synergy between two drugs. **h-j**, HT-144 cells were transfected with ASO-4 (5 nM) and then treated with vemurafenib (100 nM) in 1% ITS medium for one day. RT-qPCR assay analyzed DNFA gene expression from treatment groups relative to expression under ASO-Neg and DMSO treatment at day 1 (normalized as 1). Data are presented as mean ± SD and quantified from triplicates. One-way ANOVA tests were performed. ns, not significant; *, P < 0.05; **, P < 0.01; ***, P < 0.001; ****, P < 0.0001. **k**, HT-144 cells were transfected with ASO-4, and then treated in combination with vemurafenib. Cell viability was measured by cell titer-glo assay three days after combined treatment in 1% ITS medium. **l**, the drug interaction in **k** was evaluated by Bliss independence model.

To explore the mechanistic role of DNFA in vemurafenib resistance, we investigated immediate impact of vemurafenib on DNFA gene expression by performing time course assays on HT-144 cells in 1% ITS culture medium. After one-day treatment, vemurafenib exerts low-dose stimulation and high-dose inhibition of DNFA gene expression, exhibiting hormesis^67^ (i.e. bell-shaped dose-response curves; Supplementary Fig. 10a-g). We observed a dose-dependent induction of *PPARGC1a* expression with vemurafenib treatment (Supplementary Fig. 10h) as expected, given that the BRAF/MEK pathway directly suppresses *PPARGC1a* expression and fatty acid oxidation in melanomas^68,69^. DNFA inhibition in response to high-dose vemurafenib treatment may correlate with onset of cell death, whereas DNFA stimulation at low-dose vemurafenib treatment suggests rapid cellular resistance response. We observed that *SREBF1* depletion by ASO-4 abolished the *SCD* and *ASCL1* induction accompanying low-dose vemurafenib treatment (Fig. 4h-j). This confirms that DNFA stimulation by vemurafenib depends on SREBP1.

Consistently, vemurafenib exerts little induction of DNFA gene expression and no dose-dependent induction of *PPARGC1a* expression in A375 cells (Supplementary Fig. 11a-h), a cell line with reported MEK/ERK reactivation associated with vemurafenib treatment^70^. After treatment with an ERK inhibitor together with vemurafenib to achieve complete inhibition of BRAF/MEK/ERK pathway^71^, similarly to current clinical practice^72^, we observed strong induction of DNFA gene expression (Supplementary Fig. 11i). Our overall interpretation is that BRAF/MEK/ERK pathway inhibition promotes elevated DNFA gene expression, which then contributes to vemurafenib tolerance in melanoma cells.

Finally, we investigated treatment with ASO-4 in combination with vemurafenib. We found that that ASO-4, whether alone or in combination with vemurafenib, effectively killed HT-144 cells (Fig. 4k) and A375 cells (Supplementary Fig. 11j). However, there is a mild antagonistic effect between ASO-4 and vemurafenib at low doses by Bliss analysis (Fig. 4l and Supplementary Fig. 11k). We regard this as corollary to DNFA stimulation by vemurafenib (Fig. 4h-j and Supplementary Fig. 10a-g), while noting that, consistent with our findings above for A375, vemurafenib treatment alone yielded little induction of DNFA gene expression in A375 compared to HT-144, and the antagonistic effect was correspondingly lower. In both cell lines, high-dose ASO-4 exerts dominant cell killing effect over vemurafenib in combination treatment.

## Discussion

Cancers frequently exhibit reprogrammed metabolic traits such as elevated DNFA^3^ that act to sustain active proliferation and cell survival under adverse conditions, and support the process of tumorigenesis and metastasis, as well as resistance to targeted therapies. The diverse genetic paths cancers take to achieve such traits have frustrated efforts to exploit them clinically^73–75^.

Most DNFA enzymes are primarily regulated at the transcriptional level in a coordinated manner^76^, thus mRNA abundance of DNFA genes can be employed as a simple surrogate for DNFA activities. We find that elevated expression of multiple DNFA genes is prevalent in many cancers, suggesting that cancer cells depend upon high DNFA. Our results further show that elevated mRNA expression of DNFA enzymes (e.g. SCD) may serve as a prognostic marker for some cancer types, even though they are not considered onco-drivers. Moreover, the DNFA pathway appears to be linked to malignant cancer cell types independently of cancer-associated onco-drivers including constitutively active *BRAF* mutants. Our data suggest that elevated DNFA gene expression is intrinsic to malignant cells in melanomas, regardless of proliferative state and oncogenic mutation status. This may be consistent with the notion of increased DNFA as an “oncosustenance” pathway, with well-definable mechanistic contributions to cancer survival and proliferation. Once the oncosustenance pathway becomes active, it may not always matter which onco-drivers acted prior to cancer onset. Indeed, it has already been suggested that malignant cancer cells may have ongoing reliance upon oncogene-induced signaling pathways, but not upon initial onco-drivers^77^. It is also possible that SREBP1 transcription auto-regulation^25^ might “lock in” elevated DNFA expression in malignant cells., It seems reasonable to suggest that the scope of therapeutic investigations, which frequently focus on mutated oncogenes, could be broadened to include oncosustenance mechanisms.

We demonstrated in melanomas that SREBP1 is sufficient to up-regulate DNFA genes. Inhibition of *SREBF1* by ASO treatment results in significant reduction in the expression of DNFA genes, thereby promoting cell death, and the combined effect of multiple individual DNFA enzyme inhibitors is synergistic. Previous studies have shown that SREBP1 binding and RNAP II recruitment to DNFA gene promoters represents the primary mechanism for transcription activation^78,79^. In accord with this, we observed that RNAP II accumulated at the proximal promoter regions of *SCD* and *FASN*. Under lipid-depleted (SREBP1-activating) cell culture condition, we observed two indications of productive transcription elongation. First, there was elevated RNAP II with serine 2 phosphorylation at its C-terminal domain (RNAPII-S2p)^59^ at the gene bodies of *FASN* and *SCD*. Second, we found an increase in the histone mark for transcription elongation (H3K36me3)^80^ at the gene bodies of both genes. Based on these findings, we suggest a refined mechanism for regulation of RNAP II machinery at lipogenic gene promoters as a two-step process: RNAP II recruitment to proximal promoters, followed by RNAP II release for productive elongation. This appears to explain highly active and synchronous DNFA gene expression in melanomas, and perhaps other cell types. It is currently unclear how SREBP1 triggers RNAP II release for productive elongation.

Melanomas are frequently treated with vemurafenib, a targeted therapy to inhibit the oncogenic BRAF pathway. Because it sometimes fails to inhibit MEK/ERK (key targets downstream of BRAF)^81^, current clinical regimens combine inhibitors of both BRAF and MEK for treating metastatic melanomas^72,82^. However, even when combined inhibition is achieved, resistance arises via genetic alterations that upregulate the PI3K/AKT pathway^83^. We have found that malignant melanoma cells are able to continue (and even increase) DNFA after pro-survival signaling networks are blocked (e.g. with vemurafenib). We suspect that AKT activates SREBP1 and DNFA for survival in BRAFi-resistant melanoma cells, because DNFA gene activation and elevated lipogenesis is a well-characterized output of increased PI3K/AKT signaling^84^. Regardless, we find that 1) vemurafenib treatment is associated with DNFA stimulation and 2) activation of DNFA improves viability of some melanoma cell lines both before and after they achieve resistance to combined BRAF and MEK inhibition.

Our determination that SREBP1 is mechanistically important for resistance to targeted therapies dovetails with a similar recent finding^85^, but relies on new evidence. Putting both sets of observations together, it appears that SREBP1 and its downstream DNFA targets are necessary for melanoma cell survival and drug resistance. Our findings suggest that DNFA activity may represent a potential screening tool for the development of novel melanoma therapies.

In summary, our work demonstrates that melanoma engages the DNFA pathway for cell survival and drug resistance, employing activation of SREBP1 to directly promote transcription activation and elongation of DNFA enzyme genes. This raises the intriguing notion of SREBP1 and/or DNFA enzyme inhibition for development of future melanoma therapies.

## Methods

### Cell culture and reagents

The human melanoma cell lines HT-144, MEL-JUSO, LOXIMVI, WM1552C and MeWo were kindly provided by C. Benes (MGH Center for Molecular Therapeutics). A375 cell line was purchased from American Type Culture Collection (ATCC). Maintained in a humidified incubator at 37 °C with 5% CO_2_, all cell lines were cultured in RPMI 1640 Medium (21870092, Thermo Fisher Scientific) supplemented with 10% fetal bovine serum (Gibco), plus 2 mM L-Glutamine (Gibco) and 50 U/ml Penicillin-Streptomycin (Gibco). Two types of lipid-free medium were used for assays: 0% FBS medium contained the RPMI 1640 Medium supplemented with 2 mM L-Glutamine (Gibco) and 50 U/ml Penicillin-Streptomycin (Gibco); 1% ITS medium contained the RPMI 1640 Medium supplemented with 1× Insulin-Transferrin-Selenium (ITS-G, Thermo Fisher Scientific), 2 mM L-Glutamine (Gibco) and 50 U/ml Penicillin-Streptomycin (Gibco).

The BRAF inhibitor vemurafenib (S1267) and ERK inhibitor SCH772984 (S7101) were purchased from Selleck Chemicals. SCD inhibitor MF-438 (569406, Sigma) and FASN inhibitor GSK 2194069 (5303, Tocris) were dissolved in dimethyl sulfoxide (DMSO) to yield 50 mM stock solutions for *in vitro* studies. To generate vemurafenib-resistant cells, parental cells were exposed to increasing concentrations of vemurafenib (from 1 μM to 2 μM) for three months. The resistance was confirmed by measuring cell viability under vemurafenib treatment.

### TCGA data analysis

We analyzed 10,210 TCGA samples from 30 cancer types, for which RNA-Seq data were publically available. Briefly, gene-level RNA-Seq expression data (normalized RSEM (RNA-seq by expectation-maximization) value) were obtained from cBioportal (http://www.cbioportal.org). For Kaplan-Meier plots, RNA-Seq expression data and patient survival data from TCGA all cancers data set (10,210 samples) or TCGA skin cutaneous melanoma (SKCM) data set (476 samples) were obtained from UCSC Xena (https://xenabrowser.net). To compare the gene expression in normal skin tissues and skin tumors (Fig S2b, c), we used the uniformly analyzed RNA-Seq expression data of TCGA and GTEx (analyzed by TOIL method^86^) from UCSC Xena. The principal component analysis (PCA) in Fig S2c was performed with the R function prcomp. To compare the gene expression with oncogenic mutations in TCGA skin tumor study groups(Supplementary Fig. 4), RNA-Seq and oncogene mutation data were obtained and analyzed using CGDS R package^87^.

### Plasmids, siRNAs and ASOs

pcDNA3-Flag-nSREBP1a (plasmid #26801) and pcDNA3-Flag-nSREBP1c (plasmid #26802) were purchased from Addgene^55^. HT-144 cells were transfected with plasmids with Lipofectamine 2000 reagent (Thermo Fisher Scientific). The human-specific siRNAs targeting *SREBF1* (6720), *SREBF2* (6721), *MED15* (51586) and *CREBBP* (1387) were pre-designed ON-TARGETplus SMARTpool siRNA reagents from Dharmacon. Each ON-TARGETplus SMARTpool siRNA was a mixture of four siRNA duplexes. Sequences of individual siRNAs in each SMARTpool reagent were as follows:

siSREBF1-05 (J-006891-05): 5’-GCGCACUGCUGUCCACAAA-3’
siSREBF1-06 (J-006891-06): 5’-GAAUAAAUCUGCUGUCUUG-3’
siSREBF1-07 (J-006891-07): 5’-CGGAGAAGCUGCCUAUCAA-3’
siSREBF1-08 (J-006891-08): 5’-GCAACACAGCAACCAGAAA-3’
siSREBF2-05 (J-009549-05): 5’-GGACAGCGCUCUGGCAAAA-3’
siSREBF2-06 (J-009549-06): 5’-GCACACUGGUUGAGAUCCA-3’
siSREBF2-07 (J-009549-07): 5’-GCAGUGUCCUGUCAUUCGA-3’
siSREBF2-08 (J-009549-08): 5’-GAAAGGCGGACAACCCAUA-3’
siMed15-09 (J-017015-09): 5’-CCAAGACCCGGGACGAAUA-3’
siMed15-10 (J-017015-10): 5’-GGGUGUUGUUAGAGCGUCU-3’
siMed15-11 (J-017015-11): 5’-GGUCAGUCAAAUCGAGGAU-3’
siMed15-12 (J-017015-12): 5’-CCGGACAAGCACUCGGUCA-3’
siCREBBP-06 (J-003477-06): 5’-GCACAGCCGUUUACCAUGA-3’
siCREBBP-07 (J-003477-07): 5’-UCACCAACGUGCCAAAUAU-3’
siCREBBP-08 (J-003477-08): 5’-GGGAUGAAGUCACGGUUUG-3’
siCREBBP-09 (J-003477-09): 5’-AAUAGUAACUCUGGCCAUA-3’

The negative control siRNAs were the ON-TARGETplus Non-Targeting Pool reagents (D-001810-10, Dharmacon). Single-stranded antisense oligonucleotides (ASOs) for SREBP1 were designed and purchased as the locked nucleic acid (LNA) gapmers with phosphorothioate bonds from Exiqon. Sequences of individual ASOs were as follows:

ASO-Neg: 5’− +C*+G*+A*A*T*A*G*T*T*A*G*T*A*+G*+C*+G − 3’
ASO-1: 5’− +G*+C*+G*C*A*A*G*A*C*A*G*C*+A*+G*+A*+T − 3’
ASO-2: 5’− +T*+A*+A*G*G*G*G*A*G*T*T*A*+A*+C*+G*+G − 3’
ASO-3: 5’− +T*+A*A*G*G*T*T*T*A*G*A*G*+G*+G*+T*+G − 3’
ASO-4: 5’− +C*+T*+T*+A*G*G*G*T*C*A*A*G*A*T*+C*+G − 3’
ASO-5: 5’− +C*+A*+G*C*A*G*A*T*T*T*A*T*+T*+C*+A*+G − 3’
ASO-6: 5’− +A*+C*+C*+G*T*A*G*A*C*A*A*A*+G*+A*+G*+A − 3’

+ indicates a 2’-O, 4’-C-methylene-linked bicyclic ribonucleoside (LNA). * indicates a phosphorothioate internucleotide linkage.

siRNAs were suspended in RNase-free 1× siRNA Buffer (Dharmacon) to yield 20 μM stock solutions. ASOs were suspended in sterile ddH_2_O for stock solutions. Melanoma cell lines were transfected with siRNAs or ASOs using Lipofectamine RNAiMAX transfection reagent (Thermo Fisher Scientific) and the reverse transfection protocol suggested by the manufacture. Cells were subjected to immunoblotting or RT-qPCR analyses three days after transfection.

### Reverse transcription quantitative PCR (RT-qPCR) and RNA-Seq assays

mRNA was isolated from cultured cells using the RNeasy Mini Kit (Qiagen). RNA was treated with RNase-free DNase (Qiagen). RNA concentrations were quantified with Qubit™ RNA BR Assay Kit (Thermo Fisher Scientific). One μg RNA was used for cDNA synthesis with RNA to cDNA EcoDry™ Premix (TaKaRa) containing both random hexamer and oligo(dT)_18_ primers (Double Primed). qPCR was carried out in triplicates on a LightCycler® 480 (Roche) using LightCycler® 480 SYBR Green I Master (Roche). qPCR primers were designed by MGH primer bank (https://pga.mgh.harvard.edu/primerbank/) and the primer sequences are listed in table 1. Relative gene expression levels were calculated using the 2^−ΔΔCt^ method^88^, normalized to the 18S housekeeping gene, and the mean of negative control samples was set to 1.

**Table 1.**
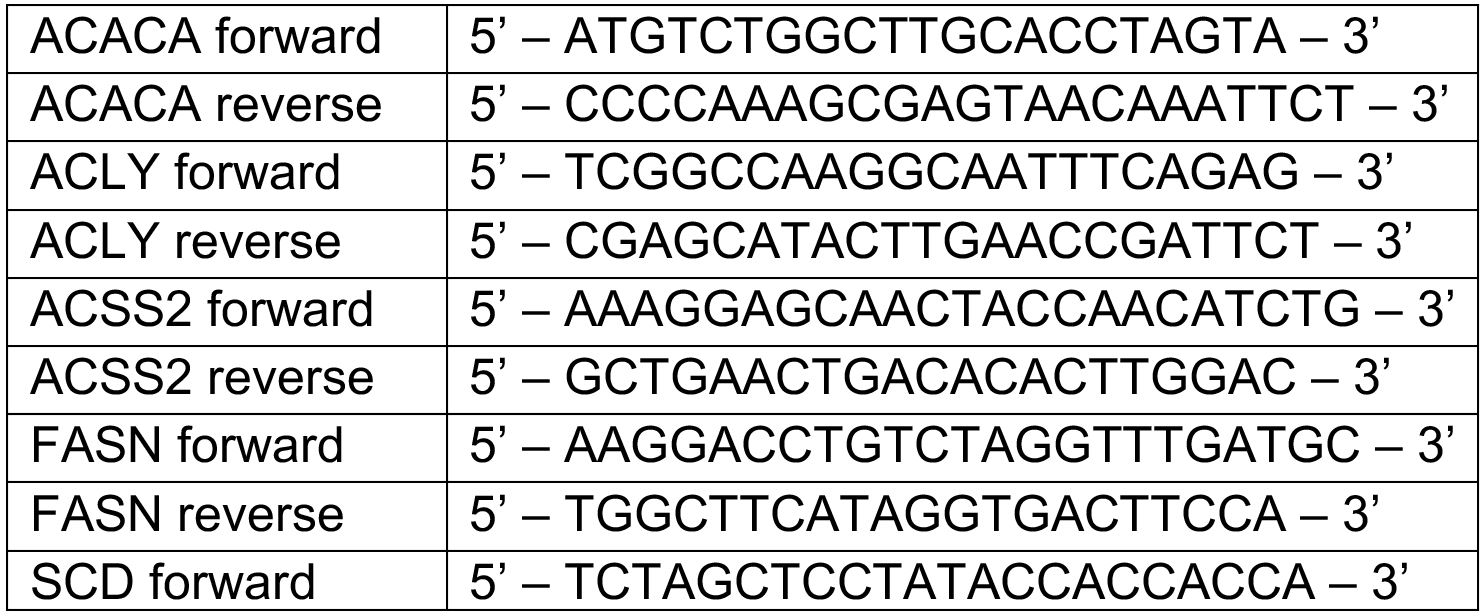
The following primers were used for RT-qPCR

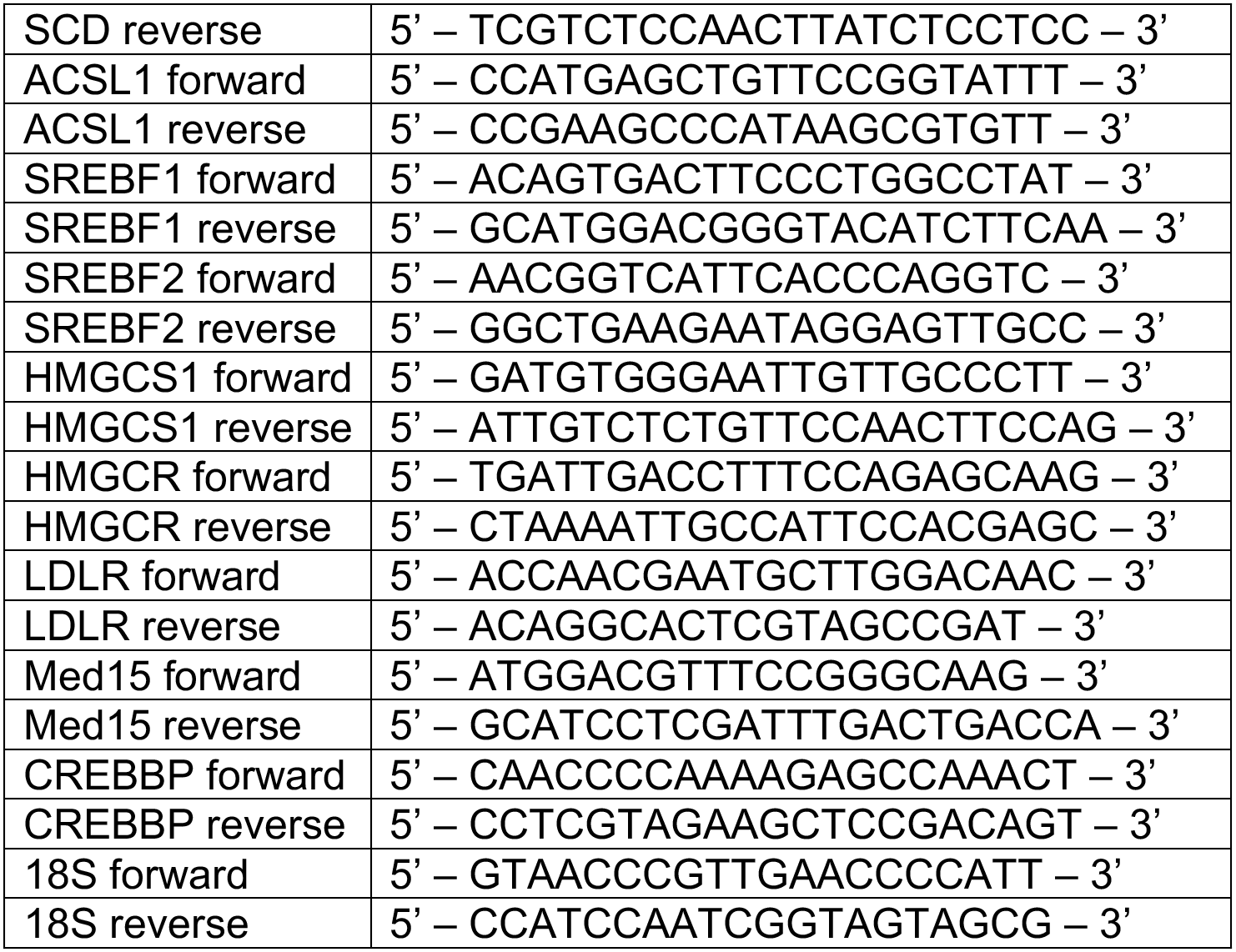

For RNA-Seq, RNA samples were harvested and treated with RNase-free DNase (Qiagen). RNA quality was verified with the RNA ScreenTape (Agilent) on Agilent 2200 TapeStation (Agilent). mRNAs were isolated from total RNA for RNA library preparation using NEBNext® Poly(A) mRNA Magnetic Isolation Module (NEB). RNA-Seq libraries were generated using NEBNext® Ultra™ Directional RNA Library Prep Kit for Illumina® (NEB) and quantified using KAPA Library Quantification Kit (KAPABiosystem). Adaptor indexed strand-specific RNA-Seq libraries were pooled and sequenced using the pair-end 75-bp per read setting for NextSeq® 500 High Output v2 Kit (150 cycles) on the NextSeq® 500 sequencer (Illumina)

### Immunoblotting assay

Total cell lysate was prepared with RIPA buffer with protease inhibitors (Protease Inhibitor Cocktail Tablets, Roche). Nuclear and cytoplasmic protein fractions were prepared with NE-PER™ Nuclear and Cytoplasmic Extraction Reagents (Thermo Fisher Scientific). Protein samples were separated on the SDS-PAGE gels using 4-15% Mini-PROTEAN® TGX™ Precast Gels (Bio-Rad) and then transferred to polyvinyl difluoride (PVDF) membranes (Immobilon-P, Millipore) for immunoblotting analysis. The following primary antibodies were used: mouse anti-SREBP-1 (IgG-2A4, BD Biosciences), rabbit anti-FASN (C20G5, Cell Signaling), rabbit anti-SCD (23393-1-AP, Proteintech), rabbit anti-ACSL1 (D2H5, Cell Signaling), rabbit anti-ACSS2 (D19C6, Cell Signaling), rabbit anti-histone H3 (9715, Cell Signaling) and rabbit anti-actin (13E5, Cell Signaling). After being incubated with primary antibodies overnight in PBST solution with 5% non-fat dry milk, membranes were probed with HRP-conjugated affinity-purified donkey anti-mouse or anti-rabbit IgG (GE Healthcare) and visualized using the Immobilon Western Chemiluminescent HRP Substrate (Millipore).

### RNA-Seq and ChlP-Seq analyses

For RNA-Seq, Illumina sequencing reads (FASTQ files) were checked with FASTQC for quality control and then aligned to the human genome (GRCh38.86). Genome index generation and sequence alignment were performed using STAR software^89^, followed by sorting and indexing of BAM files with SAMtools. Raw counts of reads mapped to genes were calculated using HT-Seq^90^. Differential expression analysis was performed using DESeq2 R package^91^. The KEGG pathway analysis of differentially expressed gene lists and principal component analysis (PCA) were performed using functions within DESeq2.

SREBP1 ChIP-Seq in multiple cancer cell lines were previously published^92^. Bam files of SREBP1 and IgG control ChIP-seq from the same cell lines were downloaded from Encode (https://www.encodeproject.org). SREBP1 binding peaks were called with Model-based Analysis of ChIP-Seq (MACS)^93^ and then annotated with ChIPseeker R package^94^. We performed *de novo* motif analysis of the SREBP1-binding sites with HOMER software (http://homer.ucsd.edu/homer/). Overlapping genes between RNA-Seq and ChIP-Seq datasets were identified by BioVenn^95^. Pathway annotation network analysis was performed on Cytoscape using ClueGo with REACTOME pathway^96^.

### ChIP quantitative PCR (ChlP-qPCR)

For each ChIP assay, 5 × 10^7^ cells were used. Chromatins from HT-144 cells were fixed with 1% formaldehyde (Polysciences) and prepared with Magna ChIP™ HiSens Chromatin Immunoprecipitation Kit (EMD Millipore). Nuclei were sonicated on a sonic dismembrator 550 (Thermo Fisher Scientific) with a microtip (model 419) from Misonix Inc. Lysates were sonicated on ice with 10 pulses of 20 sec each (magnitude setting of 3.5) and a 40-sec rest interval. The supernatant was used for immunoprecipitation with the following antibodies: rabbit anti-SREBP1 (H-160, Santa Cruz Biotechnology), rabbit anti-RNA Polymerase II (8WG16, BioLegend), rabbit anti-RNA polymerase II CTD repeat YSPTSPS (phospho S2) antibody (ab5095, Abcam), rabbit anti-RNA polymerase II CTD repeat YSPTSPS (phospho S5) (ab5131, Abcam), rabbit anti-Histone H3 (tri methyl K36) (ab9050, Abcam). qPCR reactions in triplicates were performed on a LightCycler® 480 (Roche) using LightCycler® 480 SYBR Green I Master (Roche). qPCR primers are listed in table 2.

**Table 2.**
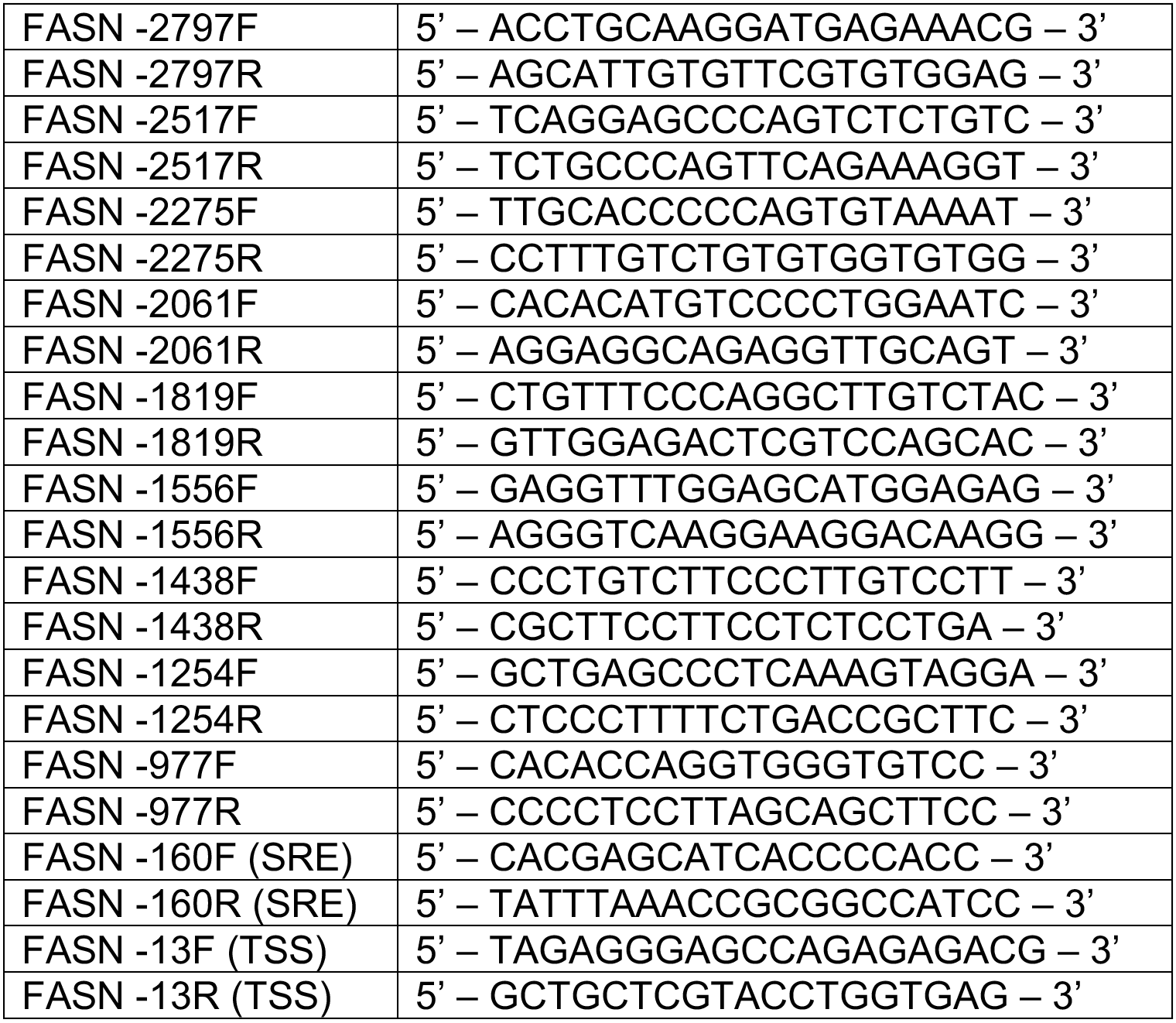
The following primers were used for ChIP-qPCR

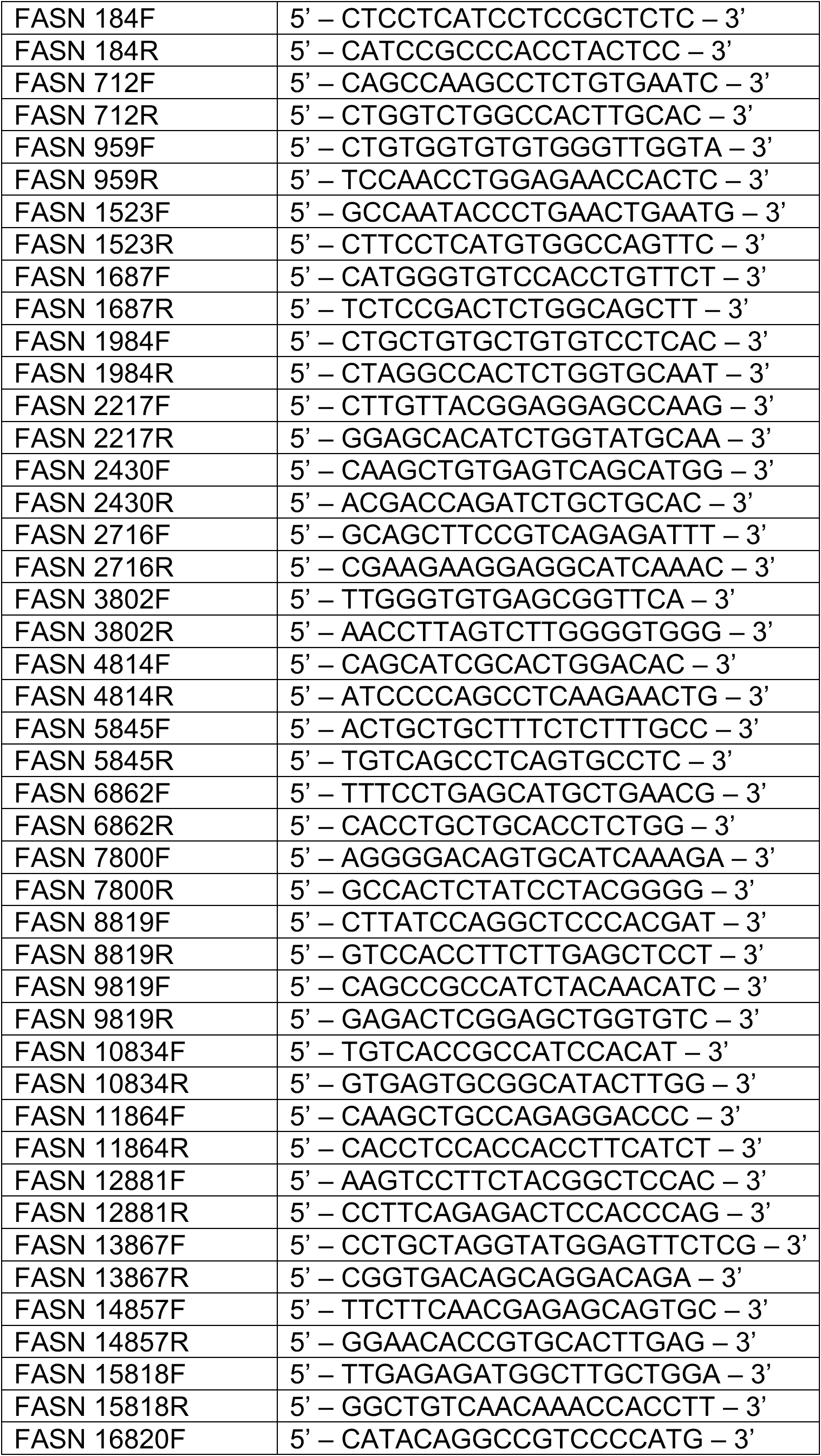

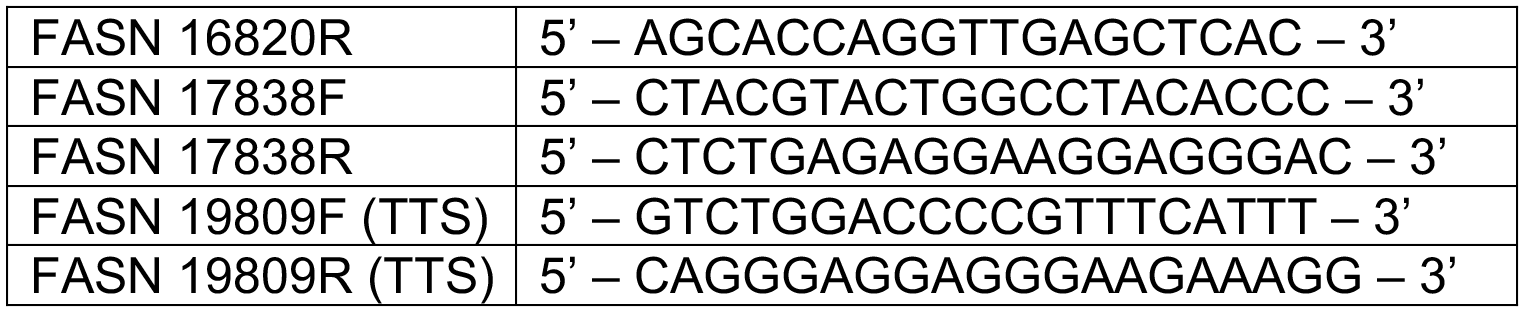

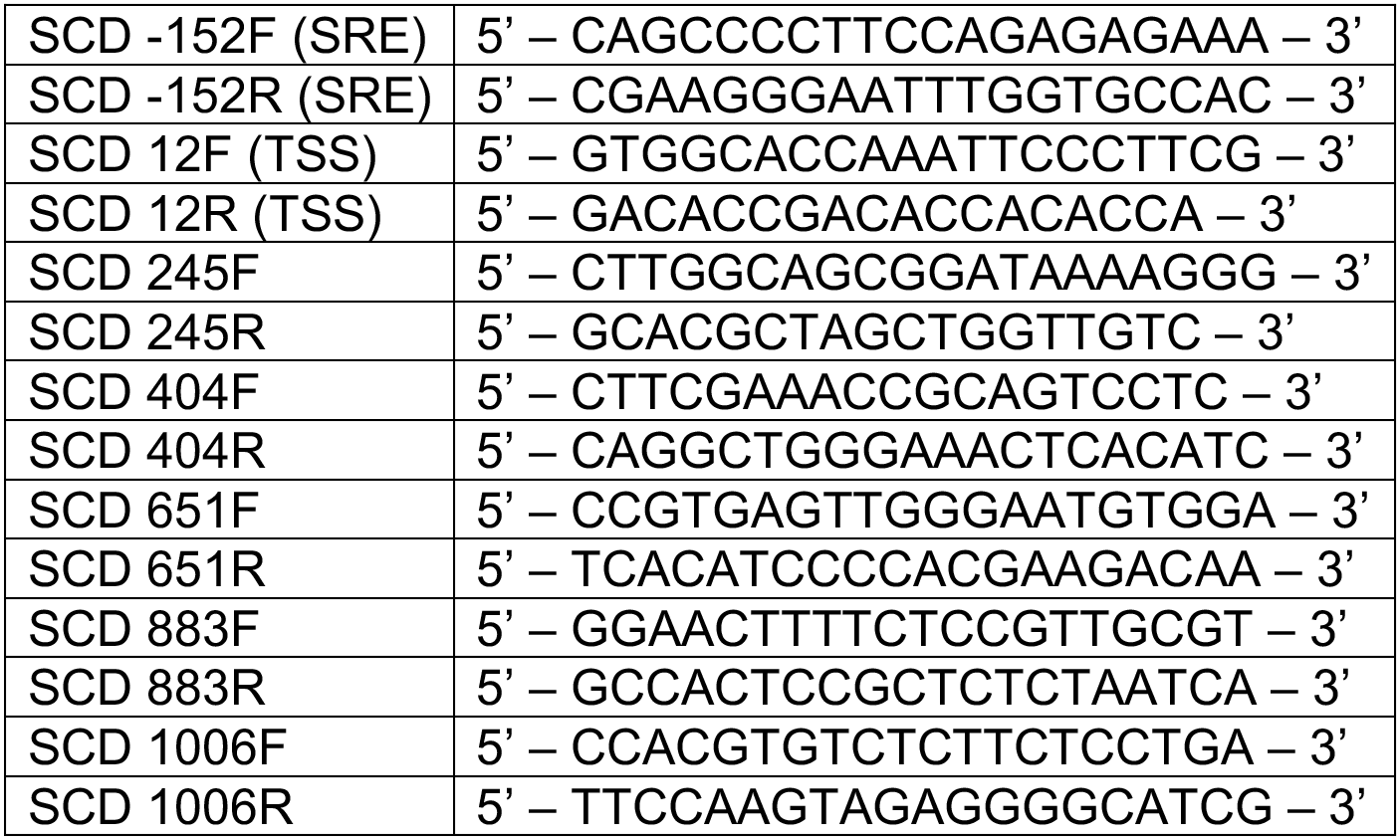

### Cell viability assay and synergy score

For dose-response assay, 1,000-2,000 cells were seeded per well in Falcon™ 96-Well Imaging Microplate with Lid (Corning). Cells were then cultured in the absence or presence of indicated drugs for 4-6 days. Relative cell viability was quantified with the CellTiter-Glo® Luminescent Cell Viability Assay Kit (G7572, Promega) and luminescent signals were measured with the Envision 2103 Multilabel Microplate Reader (Perkin Elmer). Experiments were performed independently for two or three times. Titration curves and IC50 values were generated using GraphPad Prism software. Dose-response data of drug combinations were analyzed with the SynergyFinder R package for synergism^66^. Synergy scores were calculated across all concentration combinations using Loewe or Bliss models.

## Code availability

All custom code used during the current study is available upon reasonable request.

## Acknowledgements

This work was supported by the National Institutes of Health [5R01HL116391-04] and an MGH Research Scholar Award to A.M.N. We are grateful for bioinformatics support from Drs. Andreas Petri and Sakari Kauppinen at the Aalborg University, Copenhagen, Denmark, in the design of the *SREBF1* LNA ASOs used in this publication.

## Author contributions

S.W. and A.M.N designed the study; S.W. carried out the experimental work and data analysis; S.W. and A.M.N wrote the paper. A.M.N. oversaw the project and acquired funding.

## References

1 Cairns, R. A., Harris, I. S. & Mak, T. W. Regulation of cancer cell metabolism. Nature reviews. Cancer 11, 85–95, doi:10.1038/nrc2981 (2011).

2 Vander Heiden, M. G., Cantley, L. C. & Thompson, C. B. Understanding the Warburg effect: the metabolic requirements of cell proliferation. Science 324, 1029–1033, doi:10.1126/science.1160809 (2009).

3 Vander Heiden, M. G. & DeBerardinis, R. J. Understanding the Intersections between Metabolism and Cancer Biology. Cell 168, 657–669, doi:10.1016/j.cell.2016.12.039 (2017).

4 Currie, E., Schulze, A., Zechner, R., Walther, T. C. & Farese, R. V., Jr. Cellular fatty acid metabolism and cancer. Cell Metab 18, 153–161, doi:10.1016/j.cmet.2013.05.017 (2013).

5 Menendez, J. A. & Lupu, R. Fatty acid synthase and the lipogenic phenotype in cancer pathogenesis. Nature reviews. Cancer 7, 763–777, doi:10.1038/nrc2222 (2007).

6 Brown, M. S. & Goldstein, J. L. Cholesterol feedback: from Schoenheimer’s bottle to Scap’s MELADL. Journal of lipid research 50 Suppl, S15–27, doi:10.1194/jlr.R800054-JLR200 (2009).

7 Yahagi, N. et al. A crucial role of sterol regulatory element-binding protein-1 in the regulation of lipogenic gene expression by polyunsaturated fatty acids. The Journal of biological chemistry 274, 35840–35844 (1999).

8 Xu, J., Nakamura, M. T., Cho, H. P. & Clarke, S. D. Sterol regulatory element binding protein-1 expression is suppressed by dietary polyunsaturated fatty acids. A mechanism for the coordinate suppression of lipogenic genes by polyunsaturated fats. The Journal of biological chemistry 274, 23577–23583 (1999).

9 Foretz, M. et al. ADD1/SREBP-1c is required in the activation of hepatic lipogenic gene expression by glucose. Molecular and cellular biology 19, 3760–3768 (1999).

10 Kim, J. B. & Spiegelman, B. M. ADD1/SREBP1 promotes adipocyte differentiation and gene expression linked to fatty acid metabolism. Genes Dev 10, 1096–1107 (1996).

11 Chirala, S. S. et al. Fatty acid synthesis is essential in embryonic development: fatty acid synthase null mutants and most of the heterozygotes die in utero. Proceedings of the National Academy of Sciences of the United States of America 100, 6358–6363, doi:10.1073/pnas.0931394100 (2003).

12 Horton, J. D., Goldstein, J. L. & Brown, M. S. SREBPs: activators of the complete program of cholesterol and fatty acid synthesis in the liver. The Journal of clinical investigation 109, 1125–1131, doi:10.1172/JCI15593 (2002).

13 Osborne, T. F. Sterol regulatory element-binding proteins (SREBPs): key regulators of nutritional homeostasis and insulin action. The Journal of biological chemistry 275, 32379–32382, doi:10.1074/jbc.R000017200 (2000).

14 Takeuchi, Y. et al. Polyunsaturated fatty acids selectively suppress sterol regulatory element-binding protein-1 through proteolytic processing and autoloop regulatory circuit. The Journal of biological chemistry 285, 11681–11691, doi:10.1074/jbc.M109.096107 (2010).

15 Goldstein, J. L. & Brown, M. S. 2A century of cholesterol and coronaries: from plaques to genes to statins. Cell 161, 161–172, doi:10.1016/j.cell.2015.01.036 (2015).

16 Wang, X., Sato, R., Brown, M. S., Hua, X. & Goldstein, J. L. SREBP-1, a membrane-bound transcription factor released by sterol-regulated proteolysis. Cell 77, 53–62 (1994).

17 Sato, R. et al. Assignment of the membrane attachment, DNA binding, and transcriptional activation domains of sterol regulatory element-binding protein-1 (SREBP-1). The Journal of biological chemistry 269, 17267–17273 (1994).

18 Goldstein, J. L., DeBose-Boyd, R. A. & Brown, M. S. Protein sensors for membrane sterols. Cell 124, 35–46, doi:10.1016/j.cell.2005.12.022 (2006).

19 DeBose-Boyd, R. A. et al. Transport-dependent proteolysis of SREBP: relocation of site-1 protease from Golgi to ER obviates the need for SREBP transport to Golgi. Cell 99, 703–712 (1999).

20 Sakai, J. et al. Sterol-regulated release of SREBP-2 from cell membranes requires two sequential cleavages, one within a transmembrane segment. Cell 85, 1037–1046 (1996).

21 Joseph, S. B. et al. Direct and indirect mechanisms for regulation of fatty acid synthase gene expression by liver X receptors. The Journal of biological chemistry 277, 11019–11025, doi:10.1074/jbc.M111041200 (2002).

22 Griffin, M. J., Wong, R. H., Pandya, N. & Sul, H. S. Direct interaction between USF and SREBP-1c mediates synergistic activation of the fatty-acid synthase promoter. The Journal of biological chemistry 282, 5453–5467, doi:10.1074/jbc.M610566200 (2007).

23 Reed, B. D., Charos, A. E., Szekely, A. M., Weissman, S. M. & Snyder, M. Genome-wide occupancy of SREBP1 and its partners NFY and SP1 reveals novel functional roles and combinatorial regulation of distinct classes of genes. PLoS Genet 4, e1000133, doi:10.1371/journal.pgen.1000133 (2008).

24 Oliner, J. D., Andresen, J. M., Hansen, S. K., Zhou, S. & Tjian, R. SREBP transcriptional activity is mediated through an interaction with the CREB-binding protein. Genes Dev 10, 2903–2911 (1996).

25 Amemiya-Kudo, M. et al. Promoter analysis of the mouse sterol regulatory element-binding protein-1c gene. The Journal of biological chemistry 275, 31078–31085, doi:10.1074/jbc.M005353200 (2000).

26 Mashima, T., Seimiya, H. & Tsuruo, T. De novo fatty-acid synthesis and related pathways as molecular targets for cancer therapy. British journal of cancer 100, 1369–1372, doi:10.1038/sj.bjc.6605007 (2009).

27 Ricoult, S. J., Yecies, J. L., Ben-Sahra, I. & Manning, B. D. Oncogenic PI3K and K-Ras stimulate de novo lipid synthesis through mTORC1 and SREBP. Oncogene, doi:10.1038/onc.2015.179 (2015).

28 Chen, M. et al. An aberrant SREBP-dependent lipogenic program promotes metastatic prostate cancer. Nat Genet 50, 206–218, doi:10.1038/s41588-017-0027-2 (2018).

29 Guo, D. et al. EGFR signaling through an Akt-SREBP-1-dependent, rapamycin-resistant pathway sensitizes glioblastomas to antilipogenic therapy. Sci Signal 2, ra82, doi:10.1126/scisignal.2000446 (2009).

30 Dancey, J. E., Bedard, P. L., Onetto, N. & Hudson, T. J. The genetic basis for cancer treatment decisions. Cell 148, 409–420, doi:10.1016/j.cell.2012.01.014 (2012).

31 Macconaill, L. E. & Garraway, L. A. Clinical implications of the cancer genome. J Clin Oncol 28, 5219–5228, doi:10.1200/JCO.2009.27.4944 (2010).

32 Horne, S. D., Pollick, S. A. & Heng, H. H. Evolutionary mechanism unifies the hallmarks of cancer. Int J Cancer 136, 2012–2021, doi:10.1002/ijc.29031 (2015).

33 Dienstmann, R., Rodon, J., Barretina, J. & Tabernero, J. Genomic medicine frontier in human solid tumors: prospects and challenges. J Clin Oncol 31, 1874–1884, doi:10.1200/JCO.2012.45.2268 (2013).

34 Sharma, S. V. & Settleman, J. Oncogene addiction: setting the stage for molecularly targeted cancer therapy. Genes Dev 21, 3214–3231, doi:10.1101/gad.1609907 (2007).

35 Markowitz, S. D. & Bertagnolli, M. M. Molecular origins of cancer: Molecular basis of colorectal cancer. The New England journal of medicine 361, 2449–2460, doi:10.1056/NEJMra0804588 (2009).

36 Pogrebniak, K. L. & Curtis, C. Harnessing Tumor Evolution to Circumvent Resistance. Trends Genet, doi:10.1016/j.tig.2018.05.007 (2018).

37 Salk, J. J., Fox, E. J. & Loeb, L. A. Mutational heterogeneity in human cancers: origin and consequences. Annu Rev Pathol 5, 51–75, doi:10.1146/annurev-pathol-121808-102113 (2010).

38 Lipinski, K. A. et al. Cancer Evolution and the Limits of Predictability in Precision Cancer Medicine. Trends Cancer2, 49–63, doi:10.1016/j.trecan.2015.11.003 (2016).

39 Brown, C. Targeted therapy: An elusive cancer target. Nature 537, S106–108, doi:10.1038/537S106a (2016).

40 Engelman, J. A. Targeting PI3K signalling in cancer: opportunities, challenges and limitations. Nature reviews. Cancer 9, 550–562, doi:10.1038/nrc2664 (2009).

41 Konieczkowski, D. J., Johannessen, C. M. & Garraway, L. A. A Convergence-Based Framework for Cancer Drug Resistance. Cancer Cell 33, 801–815, doi:10.1016/j.ccell.2018.03.025 (2018).

42 Rashid, A. et al. Elevated expression of fatty acid synthase and fatty acid synthetic activity in colorectal neoplasia. The American journal of pathology 150, 201–208 (1997).

43 Ide, Y. et al. Human breast cancer tissues contain abundant phosphatidylcholine(36ratio1) with high stearoyl-CoA desaturase-1 expression. PloS one 8, e61204, doi:10.1371/journal.pone.0061204 (2013).

44 Epstein, J. I., Carmichael, M. & Partin, A. W. OA-519 (fatty acid synthase) as an independent predictor of pathologic state in adenocarcinoma of the prostate. Urology 45, 81–86 (1995).

45 Porstmann, T. et al. PKB/Akt induces transcription of enzymes involved in cholesterol and fatty acid biosynthesis via activation of SREBP. Oncogene 24, 6465–6481, doi:10.1038/sj.onc.1208802 (2005).

46 Baenke, F., Peck, B., Miess, H. & Schulze, A. Hooked on fat: the role of lipid synthesis in cancer metabolism and tumour development. Dis Model Mech 6, 1353–1363, doi:10.1242/dmm.011338 (2013).

47 Bucher, N. L., Overath, P. & Lynen, F. beta-Hydroxy-beta-methyl-glutaryl coenzyme A reductase, cleavage and condensing enzymes in relation to cholesterol formation in rat liver. Biochimica et biophysica acta 40, 491–501 (1960).

48 Tirosh, I. et al. Dissecting the multicellular ecosystem of metastatic melanoma by single-cell RNA-seq. Science 352, 189–196, doi:10.1126/science.aad0501 (2016).

49 Heppt, M. V. et al. Prognostic significance of BRAF and NRAS mutations in melanoma: a German study from routine care. BMC Cancer 17, 536, doi:10.1186/s12885-017-3529-5 (2017).

50 Bennett, C. F. & Swayze, E. E. RNA targeting therapeutics: molecular mechanisms of antisense oligonucleotides as a therapeutic platform. Annual review of pharmacology and toxicology 50, 259–293, doi:10.1146/annurev.pharmtox.010909.105654 (2010).

51 Amemiya-Kudo, M. et al. Transcriptional activities of nuclear SREBP-1a,-1c, and-2 to different target promoters of lipogenic and cholesterogenic genes. Journal of lipid research 43, 1220–1235 (2002).

52 Vergnes, L. et al. SREBP-2-deficient and hypomorphic mice reveal roles for SREBP-2 in embryonic development and SREBP-1c expression. Journal of lipid research 57, 410–421, doi:10.1194/jlr.M064022 (2016).

53 Jeon, T. I. & Osborne, T. F. SREBPs: metabolic integrators in physiology and metabolism. Trends Endocrinol Metab 23, 65–72, doi:10.1016/j.tem.2011.10.004 (2012).

54 Shimomura, I., Shimano, H., Horton, J. D., Goldstein, J. L. & Brown, M. S. Differential expression of exons 1a and 1c in mRNAs for sterol regulatory element binding protein-1 in human and mouse organs and cultured cells. The Journal of clinical investigation 99, 838–845, doi:10.1172/JCI119247 (1997).

55 Toth, J. I., Datta, S., Athanikar, J. N., Freedman, L. P. & Osborne, T. F. Selective coactivator interactions in gene activation by SREBP-1a and-1c. Molecular and cellular biology 24, 8288–8300, doi:10.1128/MCB.24.18.8288-8300.2004 (2004).

56 Sedger, L. M. & McDermott, M. F. TNF and TNF-receptors: From mediators of cell death and inflammation to therapeutic giants-past, present and future. Cytokine Growth Factor Rev 25, 453–472, doi:10.1016/j.cytogfr.2014.07.016 (2014).

57 Bannister, A. J. et al. Spatial distribution of di- and tri-methyl lysine 36 of histone H3 at active genes. The Journal of biological chemistry 280, 17732–17736, doi:10.1074/jbc.M500796200 (2005).

58 Adelman, K. & Lis, J. T. Promoter-proximal pausing of RNA polymerase II: emerging roles in metazoans. Nat Rev Genet 13, 720–731, doi:10.1038/nrg3293 (2012).

59 Komarnitsky, P., Cho, E. J. & Buratowski, S. Different phosphorylated forms of RNA polymerase II and associated mRNA processing factors during transcription. Genes Dev 14, 2452–2460 (2000).

60 Liu, X., Strable, M. S. & Ntambi, J. M. Stearoyl CoA desaturase 1: role in cellular inflammation and stress. Adv Nutr 2, 15–22, doi:10.3945/an.110.000125 (2011).

61 Louie, S. M., Roberts, L. S., Mulvihill, M. M., Luo, K. & Nomura, D. K. Cancer cells incorporate and remodel exogenous palmitate into structural and oncogenic signaling lipids. Biochimica et biophysica acta 1831, 1566–1572, doi:10.1016/j.bbalip.2013.07.008 (2013).

62 Eggermont, A. M., Spatz, A. & Robert, C. Cutaneous melanoma. Lancet 383, 816–827, doi:10.1016/S0140-6736(13)60802-8 (2014).

63 von Roemeling, C. A. et al. Aberrant lipid metabolism in anaplastic thyroid carcinoma reveals stearoyl CoA desaturase 1 as a novel therapeutic target. J Clin Endocrinol Metab 100, E697–709, doi:10.1210/jc.2014-2764 (2015).

64 Jones, S. F. & Infante, J. R. Molecular Pathways: Fatty Acid Synthase. Clinical cancer research: an official journal of the American Association for Cancer Research 21, 5434–5438, doi:10.1158/1078-0432.CCR-15-0126 (2015).

65 Hardwicke, M. A. et al. A human fatty acid synthase inhibitor binds beta-ketoacyl reductase in the keto-substrate site. Nat Chem Biol 10, 774–779, doi:10.1038/nchembio.1603 (2014).

66 Aleksandr, I., He, L., Aittokallio, T. & Tang, J. SynergyFinder: a web application for analyzing drug combination dose-response matrix data. Bioinformatics, doi:10.1093/bioinformatics/btx162 (2017).

67 Reynolds, A. R. Potential relevance of bell-shaped and u-shaped dose-responses for the therapeutic targeting of angiogenesis in cancer. Dose Response 8, 253–284, doi:10.2203/dose-response.09-049.Reynolds (2010).

68 Haq, R. et al. Oncogenic BRAF regulates oxidative metabolism via PGC1alpha and MITF. Cancer Cell 23, 302–315, doi:10.1016/j.ccr.2013.02.003 (2013).

69 Vazquez, F. et al. PGC1alpha expression defines a subset of human melanoma tumors with increased mitochondrial capacity and resistance to oxidative stress. Cancer Cell 23, 287–301, doi:10.1016/j.ccr.2012.11.020 (2013).

70 Yadav, V. et al. Reactivation of mitogen-activated protein kinase (MAPK) pathway by FGF receptor 3 (FGFR3)/Ras mediates resistance to vemurafenib in human B-RAF V600E mutant melanoma. The Journal of biological chemistry 287, 28087–28098, doi:10.1074/jbc.M112.377218 (2012).

71 Kong, X. et al. Cancer drug addiction is relayed by an ERK2-dependent phenotype switch. Nature 550, 270–274, doi:10.1038/nature24037 (2017).

72 Larkin, J. et al. Combined vemurafenib and cobimetinib in BRAF-mutated melanoma. The New England journal of medicine 371, 1867–1876, doi:10.1056/NEJMoa1408868 (2014).

73 Fortunato, A. et al. Natural Selection in Cancer Biology: From Molecular Snowflakes to Trait Hallmarks. Cold Spring Harb Perspect Med 7, doi:10.1101/cshperspect.a029652 (2017).

74 Hanahan, D. & Weinberg, R. A. The hallmarks of cancer. Cell 100, 57–70 (2000).

75 Hanahan, D. & Weinberg, R. A. Hallmarks of cancer: the next generation. Cell 144, 646–674, doi:10.1016/j.cell.2011.02.013 (2011).

76 Wang, Y., Viscarra, J., Kim, S. J. & Sul, H. S. Transcriptional regulation of hepatic lipogenesis. Nature reviews. Molecular cell biology 16, 678–689, doi:10.1038/nrm4074 (2015).

77 Torti, D. & Trusolino, L. Oncogene addiction as a foundational rationale for targeted anti-cancer therapy: promises and perils. EMBO Mol Med 3, 623–636, doi:10.1002/emmm.201100176 (2011).

78 Wan, W. et al. mTORC1 Phosphorylates Acetyltransferase p300 to Regulate Autophagy and Lipogenesis. Molecular cell 68, 323–335 e326, doi:10.1016/j.molcel.2017.09.020 (2017).

79 Ponugoti, B. et al. SIRT1 deacetylates and inhibits SREBP-1C activity in regulation of hepatic lipid metabolism. The Journal of biological chemistry 285, 33959–33970, doi:10.1074/jbc.M110.122978 (2010).

80 Kolasinska-Zwierz, P. et al. Differential chromatin marking of introns and expressed exons by H3K36me3. Nat Genet 41, 376–381, doi:10.1038/ng.322 (2009).

81 Holderfield, M., Deuker, M. M., McCormick, F. & McMahon, M. Targeting RAF kinases for cancer therapy: BRAF-mutated melanoma and beyond. Nature reviews. Cancer 14, 455–467, doi:10.1038/nrc3760 (2014).

82 Robert, C. et al. Improved overall survival in melanoma with combined dabrafenib and trametinib. The New England journal of medicine 372, 30–39, doi:10.1056/NEJMoa1412690 (2015).

83 Shi, H. et al. Acquired resistance and clonal evolution in melanoma during BRAF inhibitor therapy. Cancer Discov 4, 80–93, doi:10.1158/2159-8290.CD-13-0642 (2014).

84 Krycer, J. R., Sharpe, L. J., Luu, W. & Brown, A. J. The Akt-SREBP nexus: cell signaling meets lipid metabolism. Trends Endocrinol Metab 21, 268–276, doi:10.1016/j.tem.2010.01.001 (2010).

85 Talebi, A. et al. Sustained SREBP-1-dependent lipogenesis as a key mediator of resistance to BRAF-targeted therapy. Nat Commun 9, 2500, doi:10.1038/s41467-018-04664-0 (2018).

86 Vivian, J. et al. Toil enables reproducible, open source, big biomedical data analyses. Nat Biotechnol 35, 314–316, doi:10.1038/nbt.3772 (2017).

87 Gao, J. et al. Integrative analysis of complex cancer genomics and clinical profiles using the cBioPortal. Sci Signal 6, pl1, doi:10.1126/scisignal.2004088 (2013).

88 Schmittgen, T. D. & Livak, K. J. Analyzing real-time PCR data by the comparative C(T) method. Nat Protoc 3, 1101–1108 (2008).

89 Dobin, A. et al. STAR: ultrafast universal RNA-seq aligner. Bioinformatics 29, 15-21, doi:10.1093/bioinformatics/bts635 (2013).

90 Anders, S., Pyl, P. T. & Huber, W. HTSeq--a Python framework to work with high-throughput sequencing data. Bioinformatics 31, 166–169, doi:10.1093/bioinformatics/btu638 (2015).

91 Love, M. I., Huber, W. & Anders, S. Moderated estimation of fold change and dispersion for RNA-seq data with DESeq2. Genome Biol 15, 550, doi:10.1186/s13059-014-0550-8 (2014).

92 Consortium, E. P. An integrated encyclopedia of DNA elements in the human genome. Nature 489, 57–74, doi:10.1038/nature11247 (2012).

93 Zhang, Y. et al. Model-based analysis of ChIP-Seq (mAcS). Genome Biol 9, R137, doi:10.1186/gb-2008-9-9-r137 (2008).

94 Yu, G., Wang, L. G. & He, Q. Y. ChIPseeker: an R/Bioconductor package for ChIP peak annotation, comparison and visualization. Bioinformatics 31, 2382–2383, doi:10.1093/bioinformatics/btv145 (2015).

95 Hulsen, T., de Vlieg, J. & Alkema, W. BioVenn-a web application for the comparison and visualization of biological lists using area-proportional Venn diagrams. BMC Genomics 9, 488, doi:10.1186/1471-2164-9-488 (2008).

96 Bindea, G. et al. ClueGO: a Cytoscape plug-in to decipher functionally grouped gene ontology and pathway annotation networks. Bioinformatics 25, 1091–1093, doi:10.1093/bioinformatics/btp101 (2009).

